# The Niger Delta is Under a Pollution Warning: Hydrocarbon profiles in crude oil polluted soil remediated with *Pleurotus ostreatus* and *Eisenia fitida*

**DOI:** 10.1101/2024.06.04.597352

**Authors:** Fubara Gift Evans, Dokuboba Amachree, Ilemi Jennifer Soberekon, Esther Omone Akhigbe, Diagha Opaminola Nicholas, Akayinaboderi Augustus Eli, Enyinnaya Okoro, Igoniama Esau Gamage, Ayibatonyo Markson Nathaniel, Morufu Olalekan Raimi

## Abstract

**Rationale:** The contamination of soil with crude oil poses significant environmental and ecological threats. Bioremediation, particularly through the use of organisms like *Pleurotus ostreatus* (mushroom) and *Eisenia fetida* (earthworm), has emerged as a promising approach to mitigate crude oil pollution. Understanding the effectiveness of these organisms in reducing hydrocarbon levels in contaminated soil is crucial for devising sustainable remediation strategies. **Objectives:** This study aimed to evaluate the efficacy of *Pleurotus ostreatus* and *Eisenia fetida* in remediating crude oil-polluted soil. Specifically, it sought to assess the hydrocarbon profiles in soil treated with these organisms across varying concentrations of crude oil pollution. **Method:** Crude oil concentration levels ranging from 0% to 10% were applied to soil samples, alongside control treatments including soil only, soil with earthworms, and soil with mushrooms. Each treatment was replicated five times using a randomized complete block design. Standard methods were employed to determine the hydrocarbon contents of the soil. **Results:** The results indicated a significant increase (P<0.05) in various hydrocarbon parameters, including total organic carbon (TOC), total hydrocarbons (TPH), total petroleum hydrocarbons (TPH), polycyclic aromatic hydrocarbons (PAH), and total oil and grease (TOG), with escalating concentrations of crude oil pollution. However, soil treated with *Pleurotus ostreatus* and *Eisenia fetida* exhibited noteworthy reductions in these hydrocarbon levels. At the three-month mark, mushrooms demonstrated a remarkable ability to reduce hydrocarbon content by 70%-90% compared to the pollution treatment. In contrast, earthworms exhibited minimal potential for hydrocarbon reduction, particularly at both three and six month intervals. For instance, TOC reduction reached a maximum of 96% with mushroom treatment and 85% with earthworm treatment at 5% crude oil pollution over six months. **Conclusion:** The findings highlight the effectiveness of *Pleurotus ostreatus* in significantly reducing hydrocarbon levels in crude oil-polluted soil compared to *Eisenia fetida*. Mushroom-treated soils consistently exhibited substantial reductions in TOC, TPH, TOG, PAH, and THC over the study period, suggesting their potential as a viable bioremediation agent. In contrast, while earthworms showed some capability in reducing hydrocarbon content, their effectiveness was comparatively limited. **Recommendation:** Based on the results, it is recommended to utilize *Pleurotus ostreatus* for the bioremediation of crude oil-polluted soils. Further research could explore optimizing remediation protocols involving mushroom-based treatments for enhanced efficiency. **Statement of Significance:** This study contributes valuable insights into the application of bioremediation techniques for mitigating crude oil contamination in soil. The demonstrated efficacy of *Pleurotus ostreatus* underscores its potential as a sustainable and eco-friendly solution for remediating hydrocarbon-polluted environments, offering a promising avenue for environmental restoration and conservation efforts.

## 1. Introduction

Research in environmental science, especially focusing on the Niger Delta, has thoroughly investigated the increasing rate of human influence on the wetland ecosystem of the region, reaching a point where our capacity to adequately address its repercussions is surpassed. Environmental pollution emerges as a prominent concern among the multifaceted impacts stemming from human technological progress (Abaya *et al.,* 2023a, b; Erezina *et al.,* 2023a, b; Samuel *et al.,* 2023). This pollution, stemming from various industrial activities and anthropogenic sources, exerts detrimental effects on ecosystems, endangering the well-being of humans, animals, microorganisms, and plants alike (Cook *et al.,* 2016; Morufu and Clinton, 2017; Raimi and Sabinus, 2017a, b; Olalekan *et al.,* 2017b; Raimi *et al.,* 2017; Premoboere and Raimi, 2018; Raimi *et al.,* 2018; Olalekan *et al.,* 2018a, b; Odipe *et al.,* 2018; Raimi *et al.,* 2019a, b, c, d; Suleiman *et al.,* 2019; Raimi, 2019; Olalekan *et al.,* 2019a; Henry *et al.,* 2019a, b; Olalekan *et al.,* 2020; Okoyen *et al.,* 2020; Raimi *et al.,* 2020; Deinkuro *et al.,* 2021a, b; Morufu *et al.,* 2021a, b, c; Afolabi and Morufu, 2021; Koleayo *et al.,* 2021; Olalekan *et al.,* 2021; Afolabi and Raimi, 2021; Raimi *et al.,* 2021a, b; Olalekan *et al.,* 2022a, b; Raimi *et al.,* 2022a, b, c, d, e; Raimi and Sawyerr, 2022; Omoyajowo *et al.,* 2022; Raimi and Odubo, 2022; Digha *et al.,* 2022; Ifeanyichukwu *et al.,* 2022; Clinton-Ezekwe *et al.,* 2022; Stephen *et al.,* 2022; Awogbami *et al.,* 2022; Rauf and Raimi, 2023; Kader *et al.,* 2023; Raheem *et al.,* 2023; Olalekan *et al.,* 2023; Jacob *et al.,* 2023; Raimi *et al.,* 2023; Yusuf *et al.,* 2023; Kader *et al.,* 2023a, b; Glory *et al.,* 2023; Stephen *et al.,* 2023; Sylvester *et al.,* 2023; Saliu *et al.,* 2023; Ayibatonyo *et al.,* 2024a, b; Omoyajowo *et al.,* 2024; Kwen *et al.,* 2024).

Crude oil, a natural liquid with complex hydrocarbon compositions, exemplifies one of the most significant pollutants (Sun and Li, 2013; Deinkuro *et al.,* 2021a, b; Olalekan *et al.,* 2022a; Glory *et al.,* 2023). Its pervasive contamination leads to a multitude of environmental challenges, necessitating urgent research attention. In addressing such challenges, the biological remediation approach emerges as a promising strategy for rehabilitating polluted areas and restoring ecological balance (Olalekan *et al.,* 2019b; Okoyen *et al.,* 2020; Olalekan *et al.,* 2021; Raimi *et al.,* 2022e; Sylvester *et al.,* 2023; Saliu *et al.,* 2023). Within the realm of biological remediation, fungi have garnered attention for their remarkable ability to degrade hydrocarbons, outperforming conventional methods involving bacteria (Lofthus *et al.,* 2021). Mycoremediation, a form of bioremediation employing fungi, offers a promising avenue for restoring petroleum-contaminated sites due to fungi’s capacity to detoxify pollutants and enrich soil nutrients (Adebayo *et al.,* 2021). Notably, successful trials have demonstrated the effectiveness of various fungal species, including *Pleurotus ostreatus*, *Ganoderma lucidum*, and *Trametes versicolor*, in remediating petroleum pollution. Thus, the burgeoning research landscape underscores the urgency of addressing environmental pollution and exploring innovative remediation strategies. By harnessing the potential of fungi-based mycoremediation, while we strive towards sustainable solutions for mitigating the adverse impacts of human activities on the environment.

Various species of earthworms, including *Aporrectodea tuberculata*, *Dendrobaena veneta*, *Lumbricus terrestris*, *Dendrobaena rubida*, *Eisenia fetida*, *Perionyx excavatus*, *Allobophora chlorotica*, and *Eiseniella tetraedra*, have been documented for their remarkable ability to extract contaminants such as pesticides and heavy metals from soil, including lipophilic organic micropollutants akin to polycyclic aromatic hydrocarbons (PAHs) (Liu *et al.,* 2018; Raimi *et al.,* 2018; Raimi *et al.,* 2020b; Olalekan *et al.,* 2020b; Isah *et al.,* 2020a, b; Morufu, 2021; Hussain *et al.,* 2021a, b, c; Morufu *et al.,* 2021e; Omotoso *et al.,* 2021; Asiegbu *et al.,* 2022; Modupe *et al.,* 2022a, b; Modupe *et al.,* 2023; Oshatunberu *et al.,* 2023). Additionally, earthworms play a crucial role in soil health by facilitating the transformation of approximately one quarter of organic matter into humus, with colloidal humus acting as a “slow-release fertilizer” that enriches soil fertility (Brown, 2014a, b). In recent years, there has been a growing interest in harnessing the synergistic potential of fungi and earthworms for remediating soil contaminated with crude oil. This emerging area of research holds promise as an economically viable and environmentally sustainable approach to address the challenges of crude oil pollution. Therefore, there is a pressing need to investigate the efficacy of utilizing organisms such as *Pleurotus ostreatus* and *Eisenia fetida* in reducing hydrocarbon concentrations in crude oil-contaminated soil. By exploring the remediation capabilities of these organisms, we can unlock new pathways towards mitigating the adverse impacts of petroleum pollution while promoting soil health and ecological sustainability.

## 2. Materials and Method

### 2.1 Study Area

The investigation detailed in this study was conducted within the Arboretum of Rivers State University, situated in the vibrant city of Port Harcourt, Nigeria, specifically in the area known as Nkpolu Oroworukwo. The geographical coordinates of the study site are approximately 4.7958° N latitude and 7.0246° E longitude. This region falls within the tropical wet climate zone, characterized by extended periods of heavy rainfall interspersed with brief dry spells. The average annual temperature in this locale is recorded at 26.4°C, while the average yearly rainfall amounts to 2629 millimeters, as determined by the Koppen-Geiger system (https://en.climate-data.org/).

### 2.2 Sterilization of Materials for the Experiment

Following a thorough cleaning process involving washing with detergent and subsequent rinsing with distilled water, all glassware items underwent an extensive air-drying phase. Subsequently, to ensure sterility, the glassware was individually wrapped in aluminum foil and subjected to autoclaving at a pressure of 121°C for a duration of 30 minutes. This meticulous sterilization procedure was essential to eliminate any potential contaminants and ensure the integrity of the experimental setup.

### 2.3 Source of Culture and Spawn Preparation

*Pleurotus ostreatus* cultures were sourced from Dilomat and subsequently cultivated at the Mushroom Center of Rivers State University, located in Nkpolu Oroworukwo. Here, aseptic techniques were employed to isolate and propagate pure mycelial cultures. The propagation process involved utilizing sorghum grains as the substrate for spawn production. Initially, sorghum grains underwent a meticulous preparation process. They were thoroughly rinsed and then soaked in tap water for a period of 24 hours. Following this, the soaked grains were boiled in an industrial cooker for 15 minutes, employing a 1:1 ratio of sorghum grains to water. To optimize the growth environment for the mycelium, additives such as 4% (w/w) calcium carbonate (CaCO_3_) and 2% (w/w) calcium sulfate (CaSO_4_) were incorporated, as recommended by Mohammed *et al*. (2011) and Chukunda and Simbi-Wellington (2019). Calcium carbonate served to enhance pH levels, while calcium sulfate helped prevent grain aggregation.

Once prepared, the sorghum grains were filtered to eliminate excess water and subsequently transferred into glass bottles, which were meticulously sealed using aluminum foil to maintain sterility. These bottles were then subjected to sterilization at 121°C for a duration of 30 minutes in an autoclave. After sterilization, the bottles were allowed to cool to room temperature. Subsequently, the pure *Pleurotus ostreatus* mycelial cultures were aseptically inoculated into the prepared sorghum substrate. The inoculated cultures were then incubated in darkness at a controlled temperature of 27±2°C for a period of 10-15 days. This incubation period allowed for the complete colonization of the sorghum grains by the *Pleurotus ostreatus* mycelium, ensuring robust spawn development, as described by Shyam *et al*. (2010) and Chukunda and Simbi-Wellington (2019).

### 2.4 Source and Identification of Earthworm Species

The earthworms utilized in this study were meticulously collected from the rich, moisture-laden soil of forest environments employing the meticulous hand sorting method, as outlined by Edwards (2004). Once gathered, they were carefully transferred into jars containing a portion of their native soil to maintain their natural habitat during transportation to the laboratory. Upon arrival, the earthworms underwent a comprehensive preparation process. Firstly, they were gently washed to remove any adhering soil particles, ensuring cleanliness for subsequent analysis. Following this, the earthworms were preserved in a 10% dilution concentration of formalin, a commonly used preservative solution in biological studies, to maintain their structural integrity and prevent decomposition.

To precisely identify the collected earthworm specimens, a combination of morphological and anatomical methods was employed. The morphological method described by Yousefi, Ramezani, Mohamadi, Mohammadpour, and Nemati (2009) facilitated the initial classification, while the internal anatomical method devised by Ismail (2005) provided deeper insights into their physiological characteristics. Additionally, the form and organization of setae, critical structures for earthworm locomotion and environmental interaction, were analyzed using the methodology established by Malek (2007). Among the various earthworm species identified, *Eisenia fetida* was specifically selected for its well-documented resilience to hydrocarbon exposure and other environmental stressors, as reported by Contreras-Ramos *et al*. (2008). Notably, *Eisenia fetida* is readily available and easily cultivated under laboratory conditions, making it an ideal candidate for experimental studies. As epigeic earthworms, *Eisenia fetida* primarily inhabit the upper layers of soil, where they play crucial roles in nutrient cycling and soil aeration, thus influencing ecosystem health and function.

### 2.5 Multiplication of Earthworms for Experiment

To establish a conducive environment for the identified species of earthworms, a carefully curated mixture of organic materials was prepared and utilized as a substrate for their growth and replication. This process involved a meticulous blend of cow dung, banana roots, and garden soil in a precise ratio of 3:1:1. Firstly, an appropriate area under plant shade was selected and cleared for the experimental setup. Shallow excavations, approximately 3cm deep, were then carried out using a shovel. To prevent the escape of earthworms, each excavation site was lined with a polyethylene bag placed directly on the surface, with an additional ring, such as an old motor tire, placed securely around the perimeter. Subsequently, a stratified layering approach was employed within each excavation site. Initially, clay soil was evenly spread over the polyethylene bag to absorb excess moisture and create a stable foundation. This was followed by a layer of garden soil, providing a semi-natural habitat conducive to earthworm activity. Cow dung, rich in microbial populations essential for effective organic matter degradation, was then introduced to the setup, followed by fragments of banana roots to serve as a nutrient source for the earthworms. Finally, another layer of garden soil was applied to cover the surface and provide protection.

At the center of each setup, a shallow well, approximately 5cm deep, was carefully dug. The identified earthworm specimens were then buried within these wells, ensuring their secure placement within the substrate. To maintain optimal moisture levels and prevent desiccation or escape of earthworms, the surface was thoroughly moistened with water and covered with polyethylene sheeting. Throughout the experimental period, diligent care was provided to the setups, with watering conducted twice weekly to sustain moisture levels and promote earthworm activity. This meticulous methodology was implemented over a period of three months, following guidelines provided by Anon, Biboss, Rajiv, Shristi, Sharmila, Inisa, Dhurva, and Janardan (2017), to facilitate the robust growth and replication of the earthworm population within the experimental environment.

### 2.6 Preparation of Soil for Screening for the Bioremediative Potential of *P. ostreatus* and *Eisenia fetida* on Crude Oil

A total of fifty liters of Bonny light crude oil, sourced from the Nigerian National Petroleum Corporation (N.N.P.C.) in Port Harcourt, Rivers State, Nigeria, served as the primary material for this investigation. This crude oil, renowned for its composition and relevance to regional environmental studies, formed the basis for evaluating the bioremediative potential of *Pleurotus ostreatus* and *Eisenia fetida*. The experimental methodology, adapted from Purnomo *et al*. (2010), was meticulously designed to screen the remediation capabilities of these organisms under controlled conditions. To commence the experiment, three thousand grams (3000g) of agricultural soil were precisely measured using laboratory weighing equipment and distributed into rectangular baskets covering a total area of 9.31m^2^. Each basket was lined with a cloth sack to absorb excess moisture and crude oil while simultaneously preventing the escape of earthworms. This lining facilitated adequate ventilation, crucial for the well-being of both earthworms and mushrooms throughout the experimental duration.

The soil samples were deliberately contaminated with Bonny light crude oil at varying concentrations of 5%, 7.5%, and 10%, with each treatment meticulously labeled for identification. A total of 60 baskets were utilized, with five replicates assigned to each treatment group. The experimental setup comprised two amendment treatments (mushroom and earthworm), three pollution treatments (varying crude oil concentrations), and three control treatments to ascertain the effects of amendments and crude oil pollution on soil health. The artificially polluted soil was subjected to a two-week ventilation period within the Forestry Laboratory of Rivers State University. This ventilation process, inspired by the natural weathering of crude oil in native soil, aimed to eliminate volatile toxic components from the crude oil, mimicking environmental conditions conducive to microbial activity and remediation processes (Martinkosky, 2015). Throughout this period, the contaminated soil was periodically mixed to ensure uniform distribution and exposure to ventilation.

Following the ventilation phase, 1500g of coconut coir was evenly spread over the contaminated soil and moistened with sterile distilled water. This step aimed to create an optimal growth environment for the organisms under study, ensuring suitable moisture levels for microbial and fungal activity. Subsequently, 10g of actively growing mycelium from *Pleurotus ostreatus* was aseptically inoculated into each labeled basket containing the soil-coconut coir mixture. The media were then left to incubate under ambient atmospheric conditions, monitored closely over a period of three to six months. Similarly, on the 15th day post-setup, 20g of clitelated earthworms (*Eisenia fetida*) were carefully introduced into the soil substrate. These earthworms were provided with a weekly diet consisting of a 3:1 ratio of carrot to cabbage to sustain their growth and activity. Throughout the experiment, two control groups were maintained: one comprising artificial soil without crude oil but with earthworms to assess background mortality under uncontaminated conditions, and another consisting of artificial soil with crude oil but without earthworms to monitor the natural rate of crude oil degradation in the absence of earthworm activity (adapted from Purnomo *et al.,* 2010). These control groups provided essential benchmarks for evaluating the efficacy of the experimental treatments in promoting remediation and soil health restoration.

### 2.7. Analyzing the Effects of Certain Concentrations of Crude Oil Pollution on Hydrocarbon Content of Soil Sample

Following the two-week period of soil pollution and ventilation to simulate natural weathering processes, soil samples were meticulously collected from each of the appropriately labeled pots, representing varying degrees of crude oil pollution (0%, 5%, 7.5%, and 10%). Each sample was then subjected to comprehensive analysis to determine its hydrocarbon characteristics. The analytical process involved assessing various parameters related to hydrocarbon content, including total petroleum hydrocarbon, total oil and grease, total organic carbon, polycyclic aromatic hydrocarbon, and total hydrocarbon content. These analyses were conducted in accordance with established standards outlined by the Association of Official Analytical Chemists (AOAC, 2000), ensuring methodological rigor and accuracy.

For each pollution treatment, the mean and standard error of the five replicates were meticulously calculated and reported. This statistical approach enabled a robust assessment of the impact of different concentrations of crude oil pollution on soil hydrocarbon content. By comparing these results across the pollution treatments, insights were gained into the extent of hydrocarbon contamination and the efficacy of remediation strategies. Overall, this systematic analysis provided valuable data on the effectiveness of the experimental treatments in mitigating hydrocarbon pollution and restoring soil health. These findings served as a foundation for drawing meaningful conclusions regarding the remediation potential of *Pleurotus ostreatus* and Eisenia fetida in crude oil-polluted soil environments.

### 2.8 Determining and Comparing the Effects of Bioremediators on the Hydrocarbon Content of Crude Oil Polluted Soil at Three to Six Month Intervals

To ascertain the efficacy of each bioremediation agent, namely Mushroom (represented by *Pleurotus ostreatus*) and Earthworm (represented by *Eisenia fetida*), their performance was rigorously evaluated at both three and six months within soil environments polluted with crude oil concentrations of 5%, 7.5%, and 10%. This comprehensive assessment aimed to validate their capacity to reduce hydrocarbon content in polluted soil and facilitate a comparative analysis of their remediation capabilities. To provide a baseline for comparison, the hydrocarbon content of the corresponding polluted soil samples without remediation was utilized as a reference point. The hydrocarbon parameters assessed included Total Oil and Grease (TOG), Total Petroleum Hydrocarbons (TPH), Total Hydrocarbon Content (THC), Polycyclic Aromatic Hydrocarbons (PAH), and Total Organic Carbon (TOC), with analysis conducted in accordance with established protocols outlined in the AOAC (2000) guidelines. This standardized methodology ensured the consistency and reliability of the results obtained across all testing conditions.

By examining the evolution of hydrocarbon content over the designated time intervals and across varying pollution levels, valuable insights were gained into the remediation potential of each bioremediation agent. Additionally, comparative analyses facilitated the identification of any discrepancies in performance between Mushroom and Earthworm treatments. Ultimately, this systematic evaluation served to validate the effectiveness of Mushroom and Earthworm in mitigating hydrocarbon pollution within soil environments, thus contributing to our understanding of their respective roles in environmental remediation efforts.

## Results

### 3.1 Analysis of the Effects of Certain Concentrations of Crude Oil Pollution on the Hydrocarbon Content of Soil Sample

The findings regarding the impact of varied concentrations of crude oil pollution on soil hydrocarbon content are summarized in Table 1. A notable increase in soil hydrocarbon content was observed with escalating crude oil concentrations, a trend found to be statistically significant (P<0.05). Specifically, as the concentration of crude oil in the soil increased, there was a corresponding rise in the levels of various hydrocarbon components, including Total Organic Carbon (TOC), Total Hydrocarbon Content (THC), Total Petroleum Hydrocarbon (TPH), Polycyclic Aromatic Hydrocarbon (PAH), and Total Oil and Grease (TOG). Comparatively, the soil samples with no crude oil contamination (control) exhibited the lowest levels of hydrocarbon content across all parameters. For instance, the levels of THC, TOG, PAH, TOC, and TPH in unpolluted soil ranged from 0.01, 0.1, 1.28, 2.76, to 8.98 milligrams per kilogram (mg/kg), respectively.

**Table 1:** Analyses of the Effects of Certain Concentrations of Crude Oil Pollution on the Hydrocarbon Content of Soil Sample.

| **Hydrocarbon content (mg/kg) in polluted soil sample** |  |  |  |  |  |
| --- | --- | --- | --- | --- | --- |
| **Crude Conc. (%)** | **TOC** | **THC** | **TPH** | **PAH** | **TOG** |
| 0 | 2.766 ± 0.69<sup>d</sup> | 0.01 ± 0.00<sup>d</sup> | 8.98 ± 1.15<sup>d</sup> | 1.28 ± 0.58<sup>d</sup> | 0.10 ± 0.00<sup>d</sup> |
| 5 | 5000 ± 01.15<sup>c</sup> | 4622 ± 1.15<sup>c</sup> | 4128 ± 1.15<sup>c</sup> | 874 ± 1.15<sup>c</sup> | 4698 ± 3.71<sup>c</sup> |
| 7.5 | 9870 ± 1.15<sup>b</sup> | 7792 ± 1.15<sup>b</sup> | 6348 ± 1.15<sup>b</sup> | 1974 ± 1.15<sup>b</sup> | 8401 ± 1.15<sup>b</sup> |
| 10 | 11274 ± 1.15<sup>a</sup> | 9219 ± 1.15<sup>a</sup> | 7408 ± 1.15<sup>a</sup> | 2274 ± 1.15<sup>a</sup> | 9824 ± 1.15<sup>a</sup> |

DMRT shows that the means with various letters differ considerably (P<0.05).

In contrast, following pollution with crude oil, a substantial increase in hydrocarbon content was evident. The highest mean volume of hydrocarbon was recorded at a concentration of 10% crude oil pollution, with TOC registering at 11,274 mg/kg, followed by TOG (9,824 mg/kg), THC (9,219 mg/kg), TPH (7,408 mg/kg), and PAH (2,274 mg/kg). Conversely, at a pollution concentration of 5%, the lowest hydrocarbon volumes were observed, with PAH exhibiting the least concentration at 874 mg/kg, followed by TPH (4,128 mg/kg), THC (4,622 mg/kg), TOG (4,698 mg/kg), and TOC (5,001 mg/kg). These results underscore the direct correlation between crude oil pollution levels and soil hydrocarbon content, highlighting the escalating impact of contamination on environmental integrity and necessitating effective remediation strategies. (Table 1)

### 3.2. Quantification and Comparison of the Effects of Bioremediators on Hydrocarbon Content at 0% Crude Oil Polluted Soil at Three to Six Months Interval

The outcomes of the bioremediation treatments on hydrocarbon content in soil samples with 0% crude oil pollution, assessed at three and six month intervals, are summarized in Table 2. Notably, a statistically significant decrease (P<0.05) in hydrocarbon content was observed following the application of each bioremediator (Mushroom and Earthworm) over the designated timeframes. Both Mushroom and Earthworm treatments exhibited notable reductions in hydrocarbon content, indicative of their remediation efficacy. However, the remediation treatment involving Mushroom demonstrated a more pronounced decrease in hydrocarbon levels, particularly evident at the six-month interval compared to the Earthworm treatment.

**Table 2:**
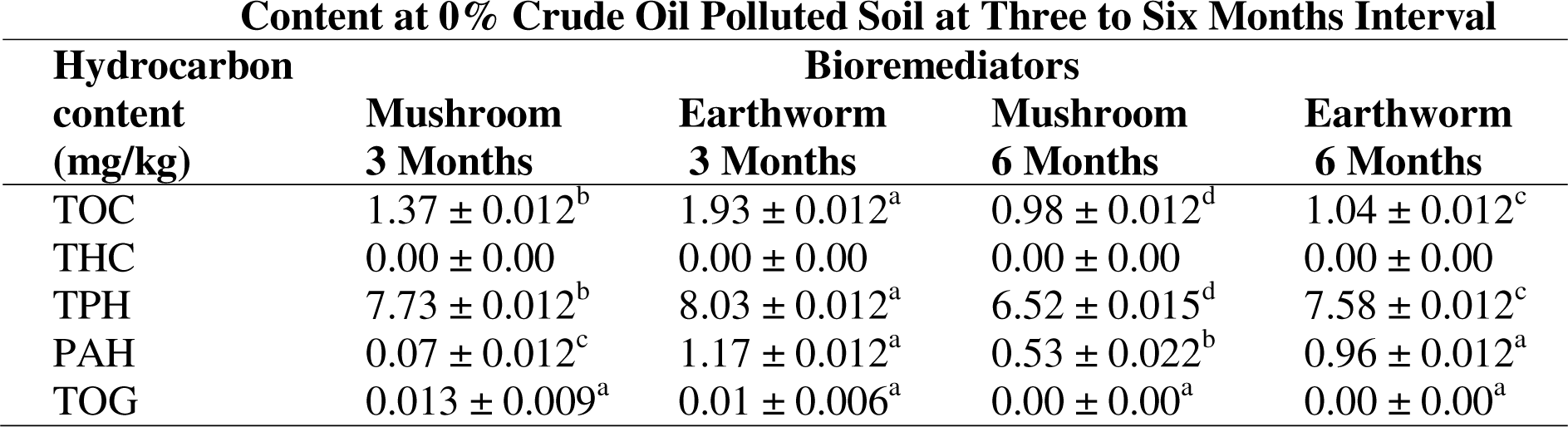
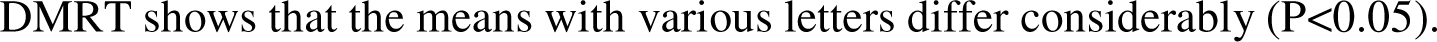
Quantification and Comparison of the Effect of Bioremediators on Hydrocarbon Content at 0% Crude Oil Polluted Soil at Three to Six Months Interval.

| Hydrocarbon content (mg/kg) | Bioremediators |  |  |  |
| --- | --- | --- | --- | --- |
|  | Mushroom 3 Months | Earthworm 3 Months | Mushroom 6 Months | Earthworm 6 Months |
| TOC | $1.37 \pm 0.012^b$ | $1.93 \pm 0.012^a$ | $0.98 \pm 0.012^d$ | $1.04 \pm 0.012^c$ |
| THC | $0.00 \pm 0.00$ | $0.00 \pm 0.00$ | $0.00 \pm 0.00$ | $0.00 \pm 0.00$ |
| TPH | $7.73 \pm 0.012^b$ | $8.03 \pm 0.012^a$ | $6.52 \pm 0.015^d$ | $7.58 \pm 0.012^c$ |
| PAH | $0.07 \pm 0.012^c$ | $1.17 \pm 0.012^a$ | $0.53 \pm 0.022^b$ | $0.96 \pm 0.012^a$ |
| TOG | $0.013 \pm 0.009^a$ | $0.01 \pm 0.006^a$ | $0.00 \pm 0.00^a$ | $0.00 \pm 0.00^a$ |

DMRT shows that the means with various letters differ considerably ( $P < 0.05$ ).

Specifically, when soil was remediated with Earthworm, the Total Petroleum Hydrocarbon (TPH) content exhibited the highest hydrocarbon levels at three months. Conversely, the Mushroom treatment induced a more substantial reduction in hydrocarbon content at six months, showcasing a 64% decrease in Total Organic Carbon (TOC) compared to 63% for Earthworms. Additionally, Mushroom treatment resulted in a 27% decrease in TPH compared to 15% for Earthworms at the six-month mark. Notably, Polycyclic Aromatic Hydrocarbon (PAH) exhibited the most significant reduction in hydrocarbon concentration, with Mushroom treatment achieving a remarkable 95% reduction compared to 25% with Earthworm treatment after six months. These findings underscore the efficacy of both Mushroom and Earthworm treatments in reducing hydrocarbon content in soil samples devoid of crude oil pollution. Moreover, the superior performance of Mushroom treatment, particularly in achieving substantial reductions in hydrocarbon content over the extended six-month period, highlights its potential as a highly effective bioremediation agent for soil remediation purposes. (Table 2).

### 3.3. Determination and Comparison of the Effects of Bioremediators on Hydrocarbon Content in Crude Oil Polluted Soil at a Three to Six Months Interval

The outcomes of bioremediation treatments on hydrocarbon content in soil samples polluted with varying concentrations (5%, 7.5%, and 10%) of crude oil, evaluated at three and six month intervals, are detailed in Tables 3a-3e. The findings reveal a statistically significant decrease (P ≤ 0.05) in hydrocarbon content following the application of bioremediators (Mushroom and Earthworm) over the specified timeframes. Overall, both Mushroom and Earthworm treatments exhibited a capacity to reduce hydrocarbon content in crude oil-polluted soil, albeit with variations in effectiveness. Notably, Mushroom treatment demonstrated a more substantial decrease in hydrocarbon content compared to Earthworm treatment, particularly evident at the six-month interval. Mushroom treatment displayed remarkable efficacy, achieving nearly 100% reduction in hydrocarbon content irrespective of the concentration of crude oil pollution at the six-month mark. Even at the three-month interval, Mushroom treatment exhibited a notable ability to reduce hydrocarbon content by 70%-90%.

In contrast, Earthworm treatment showed comparatively lower potential in reducing hydrocarbon content, with minimal improvements observed over the six-month period. While Earthworm treatment contributed to reductions in hydrocarbon content, its efficacy was notably lower compared to Mushroom treatment. Specifically, at the six-month interval, Mushroom treatment yielded impressive reductions in various hydrocarbon parameters, including TOC (up to 96%), TOG (up to 94%), TPH (up to 96.5%), PAH (up to 96%), and THC (up to 98.08%). In contrast, Earthworm treatment achieved lower reductions in hydrocarbon content, with maximum reductions ranging from 14% to 90%. These findings underscore the superior remediation potential of Mushroom treatment, particularly in achieving significant reductions in hydrocarbon content across varying concentrations of crude oil pollution. Conversely, while Earthworm treatment contributed to hydrocarbon reduction, its efficacy was comparatively limited, highlighting the differential effectiveness of bioremediation strategies in addressing soil contamination. (Tables 3a-3e).

**Table 3.**
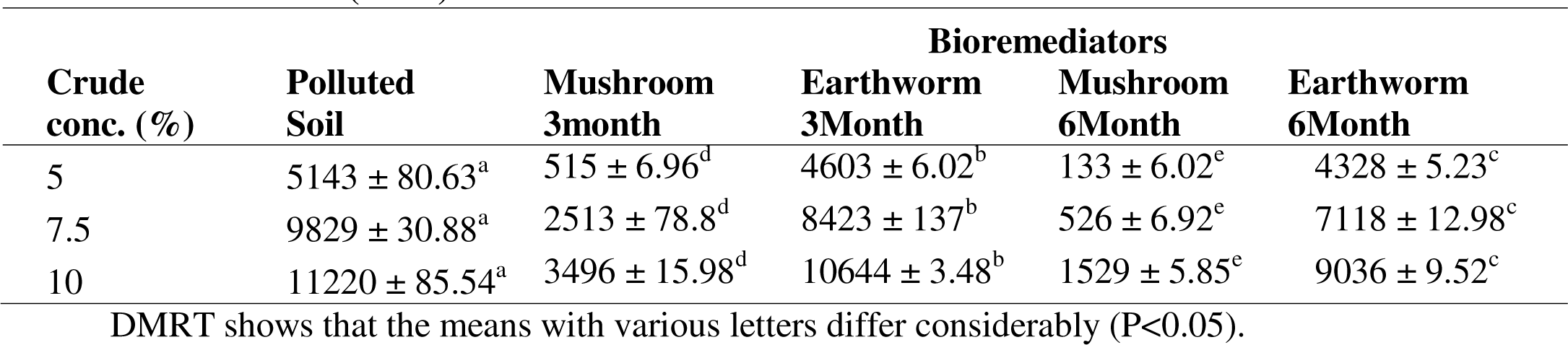
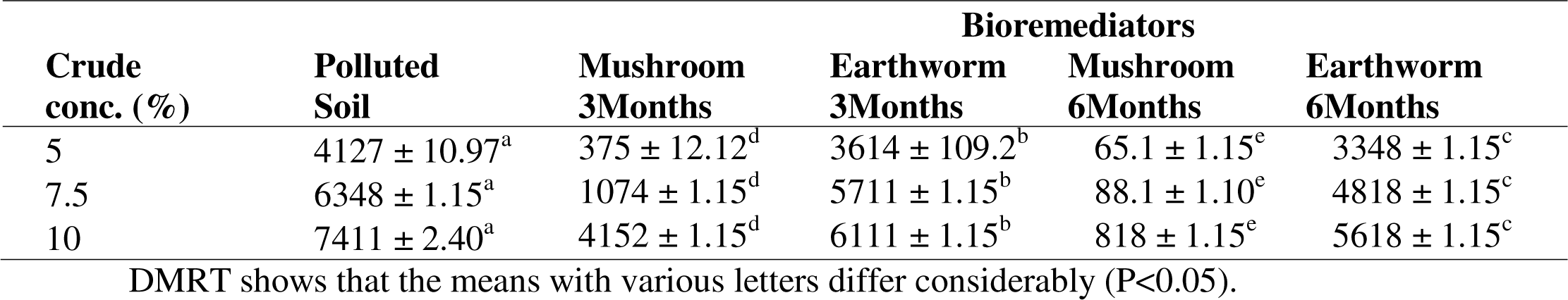
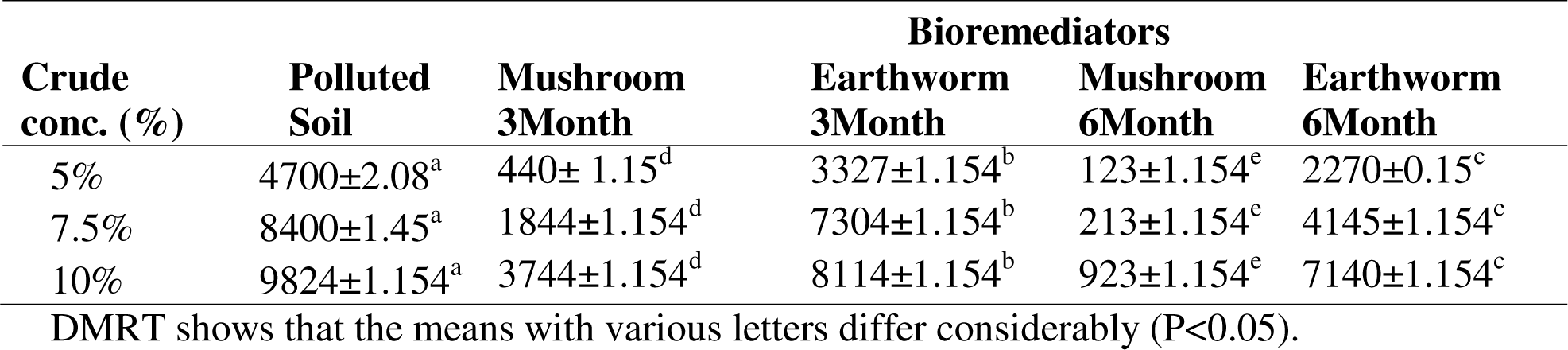
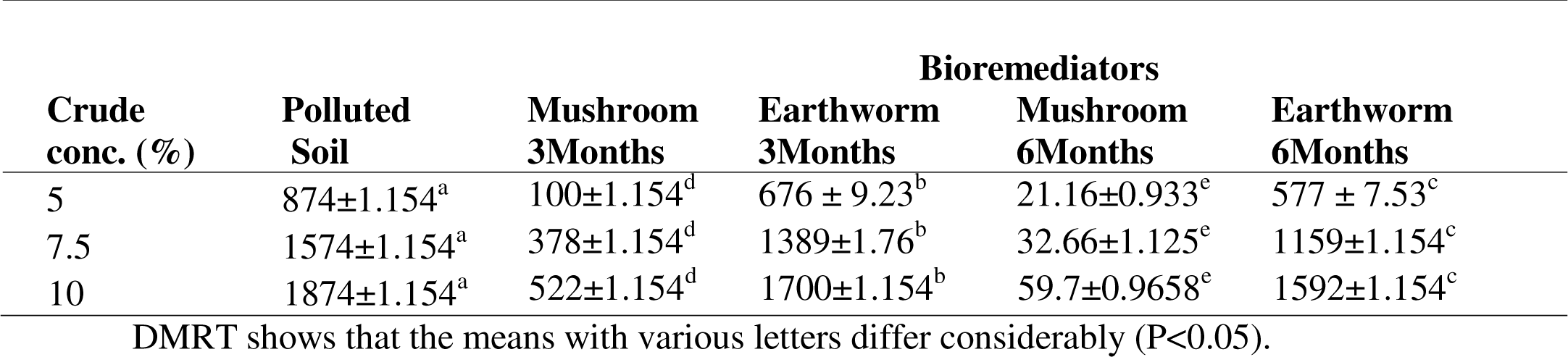
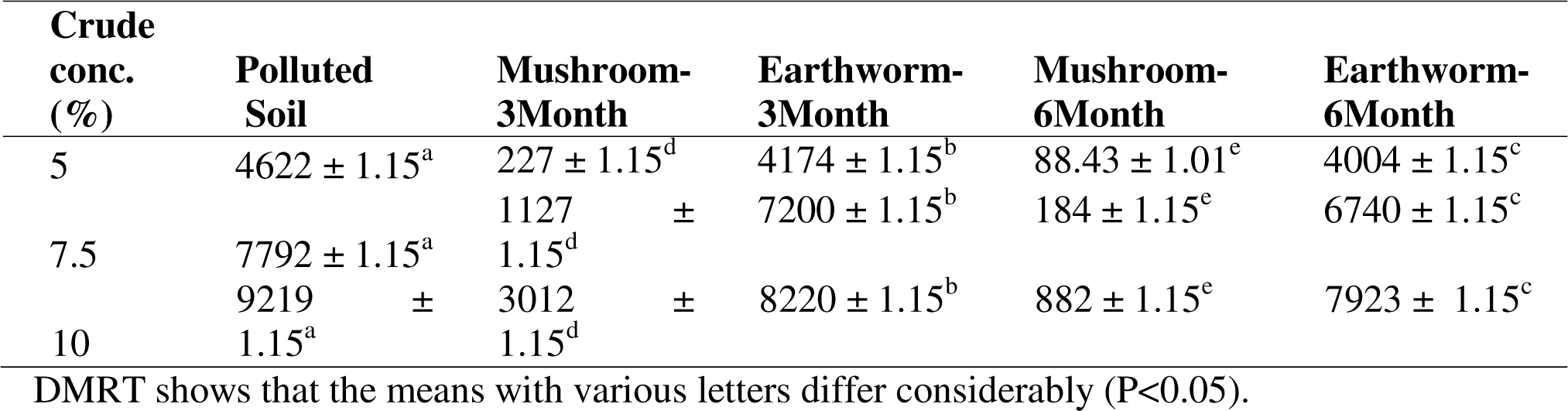
**a:** Determination and Comparison of the Effects of Bioremediators on Total Organic **b:** Determination and Comparison of the Effects of Bioremediators on Total Petroleum Hydrocarbon (TPH) in Crude Oil Polluted Soil at a Three to Six Month Interval. **c:** Determination and Comparison of the Effects of Bioremediators on Total Oil and Grease (TOG) in Crude Oil Polluted Soil at Three to Six Month **d:** Determination and Comparison of the Effects of Bioremediators on Polycyclic Aromatic Hydrocarbon (PAH) in Crude Oil Polluted Soil at a Three to Six Months Interval **e:** Determination and Comparison of the Effects of Bioremediators on Total Hydrocarbon Content (THC) in Crude Oil Polluted Soil at a Three to Six Month Interval

## 4. Discussion

### 4.1 Analysis of the Effect of Certain Concentrations of Crude Oil Pollution on Hydrocarbon Content in Soil

In this study, TOC. TPH. PAH and TOG significantly increased as the concentration of the crude oil increased in the soil sample. The hydrocarbons contents of the polluted soil with crude oil were much higher than the control. Crude oil contamination in soil influences soil enzymic activities. There is a relationship between hydrocarbons and soil enzymic activity. The degree of inhibition increases significantly with respect to increase in the levels of hydrocarbons (Alrumman *et al*., 2015). This implies that the significant high content of hydrocarbons in the crude oil polluted soil as evidenced in this study confers an inhibition of enzymic activities on soil microbes. This conclusion is comparable to the results of Vasquez-Murrieta *et al*. (2016) that PAH are ubiquitous in oil-based fuel. This is also in accordance with the findings of the report of Udoetok and Osuji (2008) that TPH, PAHs, BTEX and THC were observed at varying concentrations, even above the compliance limit in soils affected by crude oil. Osuji and Nwoye (2007) also reported presence of high extractable hydrocarbon content in crude oil contaminated soil. Zhu *et al*. (2013) observed and reported an elevation in TPH concentration in soil that had been contaminated by crude oil. The study of Wang *et al*. (2010) also corroborates the result from this study that oil contamination significantly increased TPH concentration up to 3% in relation to increased content of TOC, while studies by Alinnor *et al*., (2014) also reported that soil pollution caused by total petroleum hydrocarbons reduced as the depth increases. Furthermore, in a follow-up study by Wang *et al*. (2010), a similar finding was revealed, that the percentage of TPH in oil field marsh sampling soil was substantially greater than that of the control soil. Osuji and Onojake (2004) support the fact that high hydrocarbon thresholds impact both above and underground plants and animals that are important attributes in the cycling of nutrients that influence the accessibility of plant nutrients, in a review of various studies on the Niger Delta conducted by NDES, (1999) and Raimi and Sabinus, (2017a); Deinkuro *et al.,* (2021a, b); Ifeanyichukwu *et al.,* (2022); Olalekan *et al.,* (2022a, b); Glory *et al.,* (2023). Nevertheless, the increase in total organic carbon (TOC) in relation to the increase in crude oil pollution in our investigation does not corroborate the conclusions of Osu *et al*. (2021) who discovered a lower total organic carbon content in soil impacted by crude oil when particularly in comparison to the control treatment. They hypothesized that this was due to the oil spill altering organic carbon metabolic processes of petroleum hydrocarbons by limiting the capacity of carbon mineralization of the soil microflora. The resulting effects of soil contamination by petroleum hydrocarbons include a deterioration in the physical, chemical, and biochemical properties of the soil, a restriction in plant growth, a lack of oxygen and water, a lack of phosphorus, and nitrogen-based nutrients in the soil (Ahmed and Fakhruddin, 2018). The acidity value and heavy metal concentration levels in the soil increase due to the presence of hydrocarbon pollution or contamination (Ogboi, 2012). The contamination/pollution of the site adds several substances into the soil that differ chemically from the original composition. The lighter molecules soak through the soil and contaminate groundwater, whereas the volatile components simply evaporate (Paulauskiene *et al*., 2009). These contaminants can adhere to soil particles and stay there for a long time, while microorganisms in the soil can break down some of them through degrading mechanisms.

### 4.2 Determination and Comparison of the Effects of Bioremediators (Earthworm and Mushroom) on Hydrocarbon Content in Crude Oil Polluted Soil at Three to Six Months Interval

Various earthworm species have demonstrated efficacy in remediating herbicides, PCBs, PAHs, and other petroleum-derived hydrocarbons, as highlighted by Rafael *et al*. (2020). Building on this body of research, our study underscores the remediation potential of both earthworms and mushrooms in reducing hydrocarbon content resulting from crude oil contamination in soil. The findings reveal the remarkable ability of Mushroom and Earthworm bioremediators to significantly reduce hydrocarbon content in polluted soils. Specifically, Mushroom treatment exhibited the potential to achieve impressive reductions ranging from 94% to 96.5% in hydrocarbon content, while Earthworm treatment demonstrated a notable capability to reduce hydrocarbon content by 85% to 90% at concentrations ranging from 5% to 10% of maximum crude oil pollution over three to six months. These results underscore the promising role of both Mushroom and Earthworm bioremediation strategies in mitigating hydrocarbon pollution in soil, suggesting their potential as effective and environmentally sustainable approaches for addressing crude oil contamination and promoting soil remediation.

When comparing the decrease of all kinds of petroleum hydrocarbons from crude oil polluted soils treated with bioremediators to the control (which did not include bioremediators), it is clear that the bioremediators have the ability to eliminate hydrocarbon content from polluted soil. The mineralization of crude oil products by bioremediators (mushroom and earthworm) could be ascribed to the reduction or removal of petroleum hydrocarbons from crude oil contaminated soils, depending on the source of the petroleum hydrocarbon. This is consistent with the findings of other researchers such as Contreras-Ramos *et al*. (2008), Azizi *et al*. (2013), Tejadas and Masciandaro (2011), Raimi and Sabinus (2017a), Deinkuro *et al.,* (2021a, b) among others, who have conducted comparable study. Following 45 days of earthworm activity, Rajiv *et al*. (2013) obtained results that were consistent with those discovered in this study, including a reduction of 30 - 35 percent organic carbon and 32 - 48 percent phenol content in soil. According to Rajiv *et al*. (2013), the earthworms used in this study may have aided in the reduction of hydrocarbons out from soil via their burrowing operations, by acting as input points for nutrients and oxygen, both of which have been shown to strengthen the operations of aerobic soil microorganisms which are petroleum degraders, as well as burrowing activities that can increase the surface area for other potential degraders. Mushrooms aid in the decomposition of petroleum mostly in soil, grow in both hydrocarbon and non-hydrocarbon polluted plant substrates, and release the enzymes lignin peroxidase, manganese dependent peroxidase, and luccase all of which are employed in remediation (Mansur *et al.,* 2005). In the same vein, Stamets (2005) found that mushrooms grow best in the presence of hazardous contaminants, while Lau *et al*. (2003) found that mushroom compost may be used to decompose PAH-contaminated soil.

Barr and Aust (1994) delineated an array of white-rot fungi with the capability to degrade aromatic compounds. These fungi, specialized in lignin decomposition, exhibit exceptional prowess in modifying resilient chemical toxins such as polycyclic aromatic hydrocarbons (PAHs), as highlighted by Lang *et al*. (1995). The authors suggested that this distinctive property could be harnessed for cleansing oil-polluted soils, emphasizing the requisite of lignocellulosic substrates to support fungal survival in soil environments. Mushrooms, renowned for their diverse enzymatic capabilities, play a pivotal role in the transformation and destruction of a broad spectrum of hazardous environmental pollutants, as elucidated by Lang *et al*. (1995). Their extracellular mechanisms enable the degradation of insoluble hazardous chemicals and non-polar molecules, as noted by Levin *et al*. (2003). Additionally, mushrooms produce enzymes with low specificity, facilitating the breakdown of recalcitrant anthropogenic chemicals. In line with these observations, Amodu *et al*. (2016) demonstrated that after 50 days of cultivation, fungal (mushroom) enzymes effectively degrade various PAHs (such as pyrene, phenanthrene, and anthracene) to levels below 7% for pyrene, less than 5% for phenanthrene, and below 1% for anthracene. These findings underscore the remarkable potential of mushrooms in remediation efforts, highlighting their capacity to mitigate the impact of hazardous pollutants on the environment.

## 5. Conclusion

Crude oil comprises a complex amalgamation of hydrocarbon and non-hydrocarbon compounds, exhibiting a diverse range of toxicities to living organisms contingent upon its concentration. The repercussions of crude oil contamination on agricultural soils are profound due to its abundance of intricate molecules and persistent nature, posing challenges for effective remediation. Petroleum hydrocarbons exert detrimental effects on underground, drinking, irrigation, and industrial waters, rendering them unsuitable for various purposes. Furthermore, they pose risks to human health, the biological environment, vegetation, and can devastate natural habitats. In addressing these challenges, vermiremediation, employing earthworms to detoxify soil from environmental pollutants, and mycoremediation, utilizing fungi to mineralize, release, and sequester ions, elements, and hazardous compounds, have emerged as effective remediation techniques, as demonstrated in this study. Notably, the mushroom treatment exhibited superior remediation potential compared to earthworm-treated soils across varying levels of crude oil pollution. Consequently, this study advocates for the utilization of *Pleurotus ostreatus* and *Eisenia fitida* as efficacious agents in remediating crude oil-polluted soil.

## Authors Contribution

All authors contributed equally to conceptualization, validation, writing review and editing.

## Acknowledgments

The authors would like to express their appreciation to Dr. Morufu Olalekan Raimi as well as all anonymous reviewers, for feedback and discussions that helped to substantially improve this manuscript.

## Significance Statement

The significance of this study lies in its exploration of remediation strategies for crude oil-polluted soil, a pressing environmental concern with far-reaching implications. Crude oil contamination not only compromises soil quality but also poses risks to underground and drinking water sources, human health, biodiversity, and natural ecosystems. The complexity and persistence of petroleum hydrocarbons exacerbate the challenges of soil remediation, necessitating innovative approaches to mitigate their adverse effects. By investigating vermiremediation and mycoremediation techniques, this study offers promising solutions to address crude oil pollution in agricultural soils. The utilization of earthworms and fungi as bioremediators demonstrates their efficacy in detoxifying soil from environmental pollutants, including hydrocarbons. Furthermore, the study reveals the superiority of mushroom treatment over earthworm-treated soils in remediation effectiveness, highlighting the potential of *Pleurotus ostreatus* and *Eisenia fitida* as key agents in restoring soil health and mitigating the impacts of crude oil contamination.

Overall, the findings underscore the importance of exploring alternative remediation strategies to combat crude oil pollution effectively. The successful application of vermiremediation and mycoremediation techniques not only offers hope for restoring polluted soils but also contributes to sustainable environmental management practices. This study provides valuable insights into the remediation potential of earthworms and fungi, paving the way for future research and practical applications aimed at addressing the challenges posed by crude oil contamination in agricultural ecosystems.

## List of Abbreviation

NNPC: Nigerian National Petroleum Corporation
AOAS: Association of Official Analytical Chemists
PAH: Polycyclic Aromatic Hydrocarbons
TOG: Total Oil and Grease
TPH: Total Petroleum Hydrocarbons
THC: Total Hydrocarbon Content
TOC: Total Organic Carbon
NDES: Niger Delta Environmental Survey

